# Enhanced prefrontal nicotinic signaling as evidence of active compensation in Alzheimer’s disease models

**DOI:** 10.1101/2023.11.09.566499

**Authors:** Saige K Power, Sridevi Venkatesan, Sarah Qu, JoAnne McLaurin, Evelyn K Lambe

## Abstract

**Background:** Cognitive reserve allows for resilience to neuropathology, potentially through active compensation. Here, we examine *ex vivo* electrophysiological evidence for active compensation in Alzheimer’s disease (AD) focusing on the cholinergic innervation of layer 6 in prefrontal cortex. Cholinergic pathways are vulnerable to neuropathology in AD and its preclinical models, and their modulation of deep layer prefrontal cortex is essential for attention and executive function.

**Methods:** We functionally interrogate cholinergic modulation of prefrontal layer 6 pyramidal neurons in two preclinical models: a compound transgenic AD mouse that permits optogenetically-triggered release of endogenous acetylcholine and a transgenic AD rat that closely recapitulates the human trajectory of AD. We then tested the impact of therapeutic interventions to further amplify the compensated responses and preserve the typical kinetic profile of cholinergic signaling.

**Results:** In two AD models, we find a potentially-compensatory upregulation of functional cholinergic responses above non-transgenic controls after onset of pathology. To identify the locus of this enhanced cholinergic signal, we dissect key pre– and post-synaptic components with pharmacological strategies. We identify a significant and selective increase in post-synaptic nicotinic receptor signalling on prefrontal cortical neurons. To probe the additional impact of therapeutic intervention on the adapted circuit, we test cholinergic and nicotinic-selective pro-cognitive treatments. The inhibition of acetylcholinesterase further enhances endogenous cholinergic responses but greatly distorts their kinetics. Positive allosteric modulation of nicotinic receptors, by contrast, enhances endogenous cholinergic responses and retains their rapid kinetics.

**Conclusions:** We demonstrate that functional nicotinic upregulation occurs within the prefrontal cortex in two AD models. Promisingly, this nicotinic signal can be further enhanced while preserving its rapid kinetic signature. Taken together, our work suggests compensatory mechanisms are active within the prefrontal cortex that can be harnessed by nicotinic receptor positive allosteric modulation, highlighting a new direction for cognitive treatment in AD neuropathology.

## Introduction

Cognitive reserve denotes an individual’s ability to withstand a higher degree of neurological decline before exhibiting cognitive impairment [1]. Lifestyle factors have been shown to promote cognitive reserve, decreasing risk of dementia and slowing cognitive decline in Alzheimer’s disease (AD) [2–4]. The impact of these factors is not limited to disease onset but persists even as neuropathology worsens [5–9]. In fact, cognitive reserve is hypothesized to allow for compensation during neuropathology [10,11], generating a new or stronger response to sustain network recruitment [12,13]. In AD and its preclinical models, there are now several diverse examples of neuronal compensation leading to cognitive maintenance [14,15], but compensatory capacity in the cholinergic system has not been systematically evaluated.

Cholinergic pathways in prefrontal cortex are well positioned to contribute to mechanisms of cognitive reserve in AD [16,17]. This modulation of layer 6 of prefrontal cortex is essential for attention, executive function, and learning [18–24]. Pro-cholinergic treatments improve cognition preclinically [25–27] and slow cognitive decline clinically [28–31]. While the remodelling of the cholinergic system in AD has been extensively probed, key clinical [32–34] and preclinical studies [35–37] focus on cholinergic neurodegeneration and synapse loss. There has been relatively little functional investigation in early to mid-disease synaptic changes, except in the most swiftly-progressing genetic models of AD that preclude the use of littermate controls [38–47]. To detect and systematically investigate mechanisms of compensatory plasticity, it is essential to use a well-charted model with appropriately-matched non-transgenic controls.

To examine cholinergic signalling in AD, we use optogenetics and whole cell electrophysiology *ex vivo* to investigate prefrontal acetylcholine pathways innervating layer 6 pyramidal neurons. In models of AD from two different species, we find a strong and significant upregulation of these cholinergic responses above non-transgenic levels after onset of pathology. To probe the mechanisms underlying this functional upregulation, we systematically manipulated pre– and postsynaptic elements of the cholinergic synapse. We discovered the upregulation is specific to nicotinic receptor signalling. Lastly, we assessed the functional impact of therapeutic interventions on cholinergic responses, contrasting current standard AD treatment with a novel nicotinic treatment. Both manipulations improve endogenous cholinergic responses, but only the targeted nicotinic treatment preserves the typical rapid timing thought essential for cue detection. Overall, we demonstrate evidence for active cholinergic upregulation and put forward a mechanism of cognitive compensation that remains accessible to further potentiation with treatment.

## Materials & methods

### Animals

The University of Toronto Temerty Faculty of Medicine Animal Care Committee approved experiments in accordance with the guidelines of the Canadian Council on Animal Care (protocols #20011123, #20011621, #20011796). TgCRND8 transgenic mice expressing double human amyloid precursor protein Swedish and Indiana mutations (KM670/671NL/V717F) under the Prp gene promoter [48] on a C57BL/6 background were crossed with transgenic mice expressing channelrhodopsin in cholinergic afferents (ChATChR2; RRID:IMSR_JAX:014546, Jackson Laboratory [49]). Cholinergic neurons originating from the nucleus basalis project throughout the brain and densely innervate the deep layers of the prefrontal cortex [49,50] where we optogenetically stimulate their afferents. This transgenic model expresses channelrhodopsin under the choline acetyltransferase promoter, localizing channelrhodopsin expression to neurons that are cholinergic at the time of recording. This approach avoids expression of channelrhodopsin in neurons that are only transiently cholinergic during development [51]. TgCRND8/ChATChR2 mice were compared to ChATChR2 littermates as controls. Male and female mice were weaned at postnatal day (P)21, separated by sex, and group housed (2-4 mice per cage) in plastic cages with corn cob bedding, houses for environmental enrichment, with *ad libitum* access to food and water on a 12hr light/dark cycle with lights on at 7:00AM. Mice were used at two age groups based on previously described progression [52–56]: an early to mid-disease group and controls, 3-6 months old (n = 30, mean 4.7 ± 0.2 months), and a later disease group and controls, 7 to 12 months (n = 26, mean 9.9 ± 0.3 months). Groups were balanced for sex and genotype.

TgF344 AD rats expressing the human amyloid precursor protein Swedish mutation (KM670/671NL) and presenilin 1 with exon 9 excised, under the mouse prion promoter [57] were outbred on a Fischer strain. TgF344 AD rats were compared to F344 littermates as controls. Rats were bred at Sunnybrook Research Institute (protocol #22655), and housed in plastic cages with corn cobb bedding, 2 animals per cage, with houses and toys for environmental enrichment, on a 12hr light/dark cycle with *ad libitum* access to food and water. Rats were transferred to the University of Toronto Division of Comparative Medicine and housed for a minimum of 6 weeks before experiments. Rats were used at three age groups based on stages described previously [58–61]:, an age group at 8 months for consideration of early AD and controls (n = 21, mean 8.4 ± 0.2 months), at 12 months for mid-AD and controls (n = 23, mean 12.4 ± 0.2 months), and 18 months for later-AD and controls (n = 26, mean 17.6 ± 0.2 months). Groups were balanced for sex and genotype.

### Electrophysiology and optogenetics

Animals were anaesthetized with an intraperitoneal injection of chloral hydrate (400mg/kg) and decapitated. The brain was rapidly removed in 4°C sucrose ACSF (254 mM sucrose, 10 mM D-glucose, 26 mM NaHCO_3_, 2 mM CaCl_2_, 2 mM MgSO_4_, 3 mM KCl, and 1.25 mM Na_2_PO_4_). The 400uM thick cortical slices of the prefrontal cortex (bregma 2.2-1.1) were obtained on a Dosaka linear slicer (SciMedia). Slices were transferred to a prechamber (automate Scientific) where they recovered for at least 2 Hr in oxygenated (95% O_2_, 5% CO_2_) ACSF (128 mM NaCl, 10 mM D-glucose, 26 mM NaHCO_3_, 2 mM CaCl_2_, 2 mM MgSO_4_, 3 mM KCl, and 1.25 mM Na_2_PO_4_) at 30°C before being used for electrophysiology.

For whole-cell patch-clamp electrophysiology, brain slices were transferred to a chamber mounted on the stage of a BX51WI (Olympus) microscope and perfused with 30 °C oxygenated ACSF at 3-4mL/min. Layer 6 pyramidal neurons in the prelimbic and anterior cingulate regions of the medial prefrontal cortex were identified by size, morphology, and proximity to white matter, visualized using infrared differential interference contrast microscopy. Recording electrodes were filled with patch solution containing 120 mM K-gluconate, 5 mM MgCl_2_, 4 mM K-ATP, 0.4 mM Na_2_-GTP, 10 Mm Na_2_-phosphocreatine, and 10 mM HEPES buffer adjusted to pH 7.33 with KOH. Data were acquired and low-pass filtered at 20kHz with Axopatch 200b amplifier (Molecular Devices) and Digidata 1440 digitizer and pClamp10.3 acquisition software (Molecular Devices). Responses from a homogenous population of layer 6 regular spiking pyramidal neurons [62,63] were examined in voltage clamp at –75 mV and in current clamp.

To excite channelrhodopsin-containing cholinergic afferents optogenetically (opto-ACh), blue light (470 nm) was delivered in 5ms pulses with an LED (2mW; Thorlabs) through a 60x objective lens. This opto-ACh stimulus was delivered in a frequency-accommodating (50-10 Hz) eight pulse train to stimulate cholinergic axons [62,63] to mimic the activation pattern of cholinergic neurons [64–66]. This opto-ACh protocol elicits direct cholinergic inward current responses resilient to combined application of glutamatergic synaptic blockers [50,62,63,67] and sensitive to combined cholinergic receptor antagonists [62].

### Experimental design, analysis, and statistics

Animals of both sexes were used in pharmacological interventions. All experiments draw on multiple rodents per interventions, and recordings were made from 1 to 3 neurons per brain slice, (brain slices of prefrontal cortex in mouse: 2-3 slices; rat: 3-4 slices). Pharmacological agents were pre-applied for 10 min and co-applied during optogenetic stimuli or exogenous acetylcholine bath application and recovery period. CNQX (50 µM) and D-APV (50 µM) were applied to block glutamatergic synaptic transmission; bicuculline (3 µM) and CGP55845 (1 µM) were applied to block GABA receptors; Atropine (200 nM; Sigma-Aldrich) was applied to block muscarinic receptors; dihydro-β-erythroidine (DHβE; 3 µM) to block β2-containing nicotinic receptors; AF-DX 116 (300 nM) to block M2 muscarinic receptors; galantamine hydrobromide (1 µM) to block acetylcholinesterase, and NS9283 (1 µM) to potentiate nicotinic responses. Compounds from Tocris Biosciences unless otherwise specified.

Data analysis was performed in Clampfit 10.3 (Molecular Devices) and Axograph and statistical analysis was performed with Graphpad Prism 10. To measure fast-onset kinetics, raw traces were used for calculating rising slope of current response within 50 ms of opto-ACh onset. To measure response magnitude, downsampled traces were used to fit triple exponentials to opto-ACh responses, from which the peak amplitude (pA) and decay constant (s), where applicable, were measured.

Firing rise is measured by the mean increase in firing frequency for the initial 0.5 s following the onset of the opto-ACh stimulus relative to the mean baseline firing frequency (calculated over 2 s before opto-ACh stimulus). Peak firing frequency is measured as the difference between the mean baseline firing frequency and fastest instantaneous firing frequency following opto-ACh stimulus. Genotype differences between opto-ACh responses were assessed with unpaired two-tailed *t*-tests. Pharmacological manipulations to assess synaptic components of the cholinergic response are calculated with unpaired, two-tailed *t*-tests. The effects of therapeutic interventions are assessed with paired, two-tailed *t*-tests. The impact of sex on cholinergic measures and interactions between sex and genotype are analyzed by two-way ANOVA.

To measure currents to exogenous ACh, 1mM ACh was bath-applied for 15 s. Genotype differences in acetylcholine current magnitude were measured by peak amplitude. Acetylcholine firing response was measured by the time from stimulus onset to peak firing. Both measures are graphed with a cumulative frequency distribution with 10 pA or 10 s bin width, allowing for direct comparison of each age group’s genotype differences. Genotype differences in these measures are calculated with a nonparametric Kolmogorov Smirnov (KS) test.

Intrinsic neuronal properties are assessed in current clamp and voltage clamp recordings at baseline and statistically analysed using unpaired two-tailed *t*-tests. To measure intrinsic neuronal excitability, input-output curves are generated by measuring neuronal firing frequency in response to increasing steps of injected current. These experiments are statistically analysed by performing a second order polynomial nonlinear regression of input-output curves and a comparison of fit, analysed with an F test.

## Results

### Increased opto-ACh responses in early to mid-disease in TgCRND8 AD mouse model

To measure functional changes in prefrontal cortex cholinergic signals in an AD model, we record whole-cell responses to endogenous acetylcholine released optogenetically (opto-ACh; Fig 1A.1, A.2) in littermate non-transgenic control and TgCRND8 AD mice during early to mid-AD (3 to 6 months) [52–56]. These mice were generated by crossing TgCRND8 transgenic AD model mice [48] with ChAT-ChR2 transgenic mice expressing channelrhodopsin in cholinergic afferents [49]. Cholinergic neurons from the nucleus basalis project throughout the brain and densely innervate the deep layers of the prefrontal cortex [49,68], where they respond to optogenetic stimulation. TgCRND8/ChATChR2 mice were compared to ChATChR2 littermates as controls. Optogenetic responses were recorded from pyramidal neurons in layer 6 of the prelimbic and anterior cingulate areas of the prefrontal cortex. Layer 6 receives dense innervation of cholinergic afferents [68] and shows strong excitatory responses [69]. Responses from a homogenous population of regular spiking pyramidal neurons [62,63] were examined in voltage clamp at –75mV and in current-clamp. The brief opto-ACh protocol [62,63,67] elicits inward current responses resilient to combined application of glutamatergic synaptic blockers CNQX (20 µM) and D-APV (50 µM) in non-transgenic control (pA, opto-ACh; 39.2 ± 14 pA, opto-ACh synaptic blockers: 40.5 ± 14.3 pA; paired *t*-test: *t_5_* = 0.3, *P* = 0.8) and TgCRND8 (pA, opto-ACh; 82.7 ± 22 pA, opto-ACh synaptic blockers: 80.4 ± 19 pA; paired *t*-test: *t_4_* = 0.3, *P* = 0.8) and resilient to combined application of GABA receptor blockers BCC (3 µM) and CGP (1 µM) in non-transgenic control (pA, opto-ACh; 28.2 ± 6.5 pA, opto-ACh GABA blockers: 24.8 ± 5.9 pA; paired *t*-test: *t_5_* = 2, *P* = 0.1) and TgCRND8 (pA, opto-ACh; 45.4 ± 11.9 pA, opto-ACh synaptic blockers: 43.8 ± 11.5 pA; paired *t*-test: *t_7_* = 0.7, *P* = 0.5).

**Figure 1.**
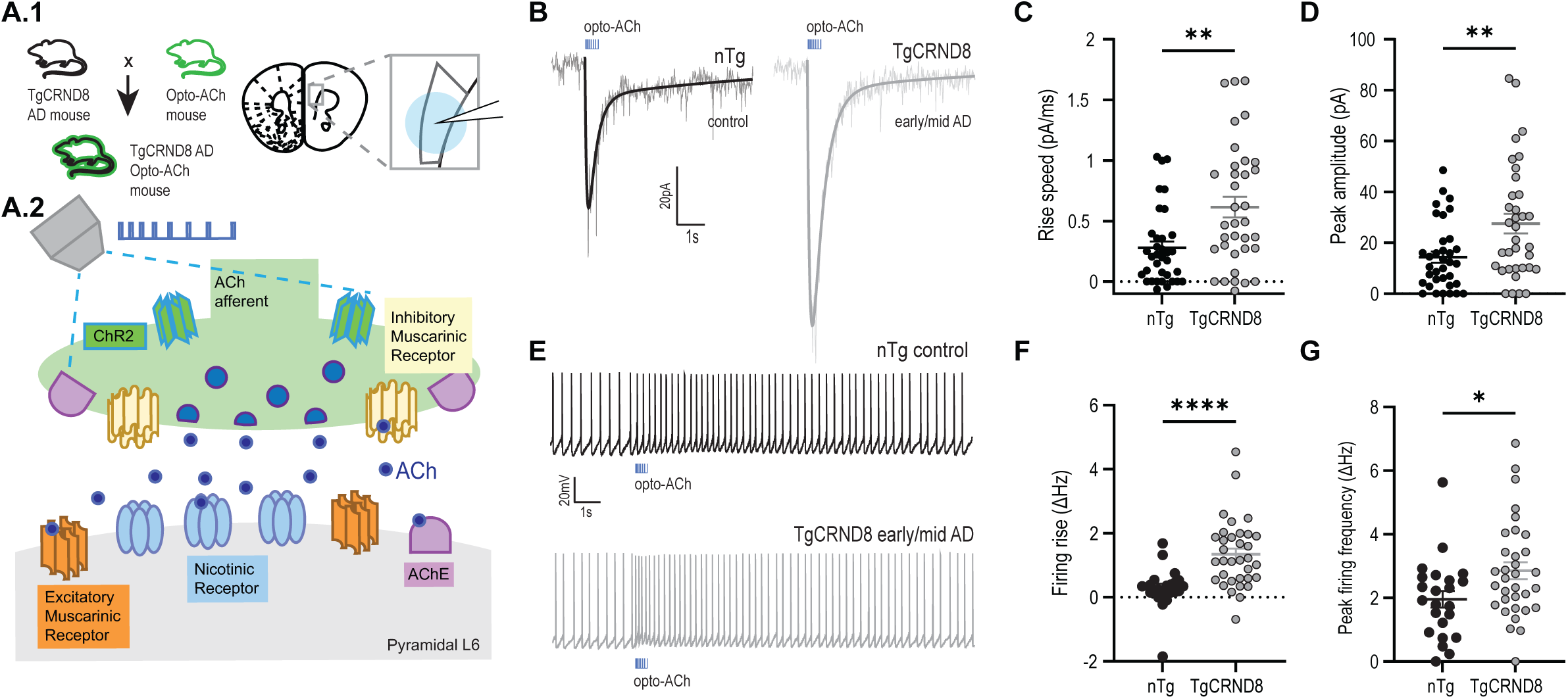
Endogenous opto-ACh responses are upregulated in early to mid-AD. **A.1** Image depicts compound transgenic mouse cross between TgCRND8 AD mouse model and ChAT-ChR2 mouse model expressing channelrhodopsin in cholinergic neurons. Image depicts coronal brain section from these mice, showing the recording electrode positioned over layer 6 of the prefrontal cortex. **A.2** Schematic adapted from Power et al., 2023, represents the cholinergic prefrontal synapse receiving optogenetic stimulus (0.5s pulse train of decreasing frequency). Afferents release endogenous acetylcholine (ACh) which binds to nicotinic and muscarinic receptors, exciting postsynaptic pyramidal neurons. **B** Example opto-ACh responses from non-transgenic (nTg) control (black) and TgCRND8 AD (grey) neurons measured in voltage clamp (Vm = –75 mV). Graphs show significant increase in **C** rise speed (*P* < 0.01) and **D** peak amplitude (*P* < 0.01) of opto-ACh currents in TgCRND8 early AD responses relative to age-matched controls (3-6 months old). **E** Example opto-ACh responses from WT control (black) and TgCRND8 AD (grey) neurons measured in current clamp. Graphs show significant increase in **F** firing rise (*P* < 0.0001) and **G** peak firing frequency (*P* < 0.05) of opto-ACh firing responses in TgCRND8 early/mid AD responses relative to age-matched controls. Mice aged 3 to 6 months (nTg: 4.5 ± 0.4 months, n = 10, TgCRND8: 4.8 ± 0.3 months, n = 10).

We find a significant increase in opto-ACh current responses in early to mid-AD mice (3 to 6 months) (Fig 1B), as measured by a significantly greater rise slope (Fig 1C, unpaired, two-tailed *t-*test: *t* = 3.3, *P* = 0.002, df = 68) and peak amplitude (Fig 1D, unpaired, two-tailed *t*-test: *t* = 3, *P* = 0.004, df = 68) in AD responses.

This upregulation is also detected in the change in the firing response elicited by opto-ACh (Fig 1E). Holding the neurons in current clamp and injecting current to elicit ∼2-3 Hz action potential firing at baseline, opto-ACh elicits a significantly greater increase in the delta initial firing frequency or firing rise (1F, unpaired, two-tailed *t*-test: *t* = 4.2, *P* < 0.0001, df = 54) and the delta peak firing frequency (1G, unpaired, two-tailed *t*-test: *t* = 2.3, *P* = 0.02, df = 54) in AD compared to control.

To delve deeper into the opto-ACh result, we investigated the effect of sex, but did not detect a main effect of this factor (two-way ANOVA, F_1, 64_ = 1.4, *P* = 0.2), nor a significant interaction between sex and genotype (F_1, 64_ = 1.8, *P* = 0.4) on cholinergic current amplitude.

The genotype differences in the opto-ACh responses in voltage-clamp and current-clamp are observed in the absence of corresponding genotype differences in intrinsic membrane properties (Supplemental Table 1). We also assess neuronal excitability by measuring action potential firing rate in response to increasing steps of current injection, generating input-output curves. In the absence of opto-ACh, we find no significant difference in the excitability curves (Fig S1A; nonlinear regression, comparison of fit, F test = ns). These findings underscore the relative specificity of opto-ACh in this AD mouse model to increase the excitability of layer 6 pyramidal neurons during early to mid-disease.

### Preserved opto-ACh responses in later-disease in TgCRND8 AD mouse model

To ask what happens to the AD cholinergic response as neuropathology progresses, we recorded from age– and sex-matched littermate non-transgenic control and TgCRND8 AD mice at a more advanced stage of AD. In this older group (7 to 12 months), opto-ACh current responses in AD mice are similar to the non-transgenic level (Fig 2A-C; unpaired, two-tailed *t*-test: rise slope *t* = 0.07, *P* = 0.9, df = 75; peak amplitude *t* = 0.5, *P* = 0.6, df = 75) as well as opto-ACh firing responses (Fig 2D-F; unpaired, two-tailed *t*-test: firing rise *t* = 0.6, *P* = 0.6, df = 60; peak firing *t* = 0.2, *P* = 0.8, df = 60). Cholinergic responses in the model are no longer significantly greater than controls at the later time point.

**Figure 2.**
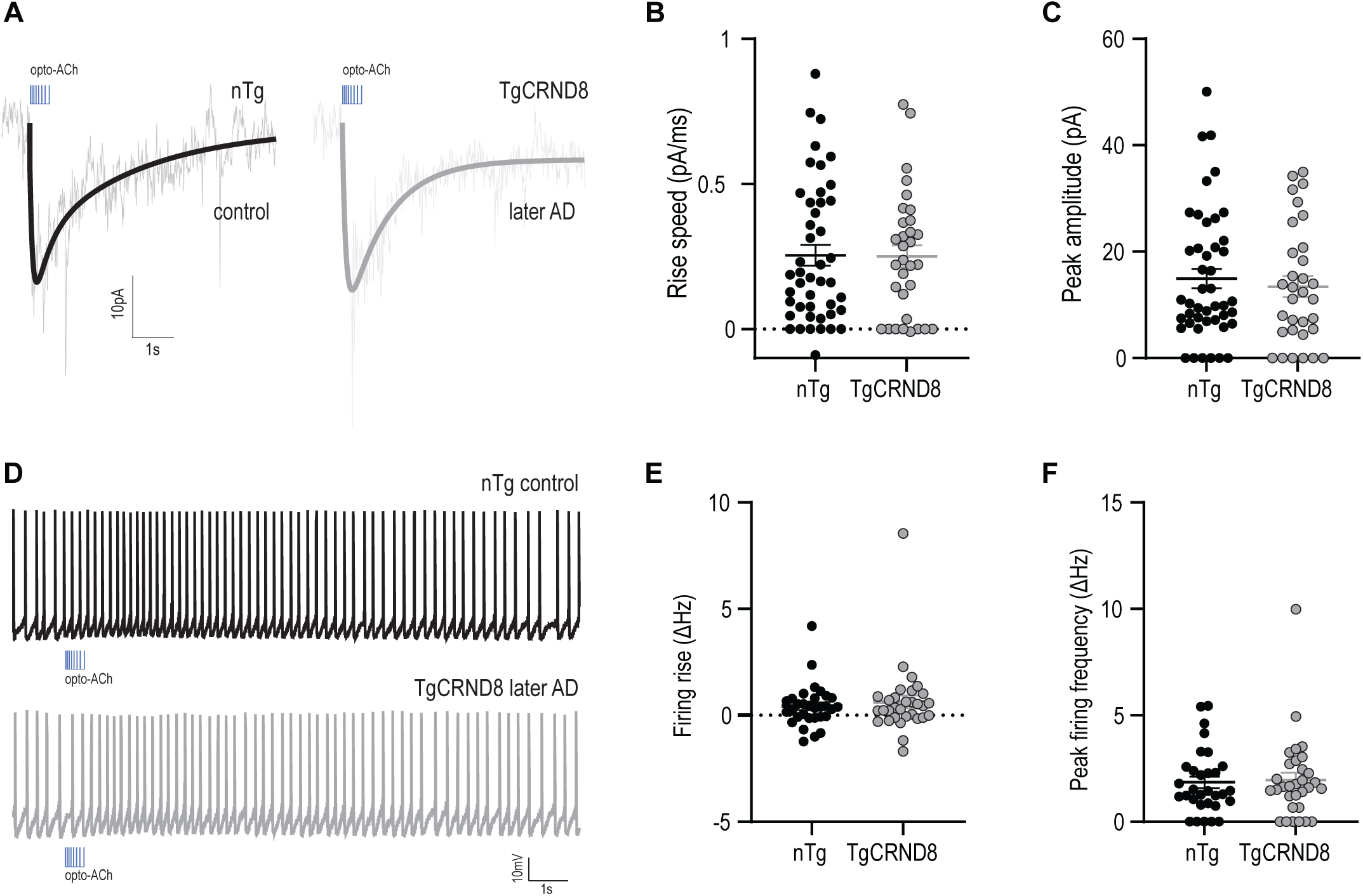
Cholinergic signaling not different from non-transgenic in late AD. **A** Example opto-ACh responses from non-transgenic (nTg) control (black) and TgCRND8 AD (grey) neurons later in disease measured in voltage clamp (Vm = –75 mV). Graphs show no difference in **B** rise speed and **C** peak amplitude of opto-ACh currents in TgCRND8 late AD responses relative to age-matched controls. **D** Example opto-ACh responses from WT control (black) and TgCRND8 AD (grey) neurons later in disease measured in current clamp. Graphs show no difference in **E** firing rise and **F** peak firing frequency of opto-ACh firing responses in TgCRND8 late AD responses relative to age-matched controls. Mice aged 7-12 months (nTg: 9.3 ± 0.5 months, n = 10, TgCRND8: 10.1 ± 0.6 months, n = 10).

Consistent with the younger age group, we find no sex differences in cholinergic responses, where there is no significant effect of sex (two-way ANOVA, F_1, 73_ = 0.1, *P* = 0.7), nor significant interaction between sex and genotype (F_1, 73_ = 2.2, *P* = 0.1) on cholinergic current amplitude.

Further investigating the intrinsic membrane properties of these neurons shows no significant genotype differences (Supplemental Table 1). Investigating neuronal excitability, input-output analysis finds no significant genotype difference in excitability curves (Fig S1B; nonlinear regression, comparison of fit, F test = ns). These results suggest that prefrontal endogenous cholinergic responses are upregulated in AD beyond WT levels in early/mid-pathology but that this high level is not maintained above non-transgenic controls when the AD model reaches later disease.

### Increased cholinergic responses at mid-disease in TgF344 AD rat model

To evaluate whether cholinergic upregulation in response to AD pathology is recapitulated across species, we consider the TgF344-AD rat model [57]. We compared TgF344-AD and littermate WT controls at three points in disease: early-AD (8 months), mid-AD (12 months), and later-AD (18 months) [58–61]. Strikingly, we again find an upregulation in cholinergic currents in TgF344-AD rats relative to controls (Fig 3A, 3B). This difference is specific to the mid-AD age group (KS test, *P* = 0.005, D = 0.4) (Fig 3C-E). It was not observed in the early-AD group (KS test, *P* = 0.7, D = 0.2), nor in the later-AD group (KS test, *P* = 0.7, D = 0.2).

**Figure 3.**
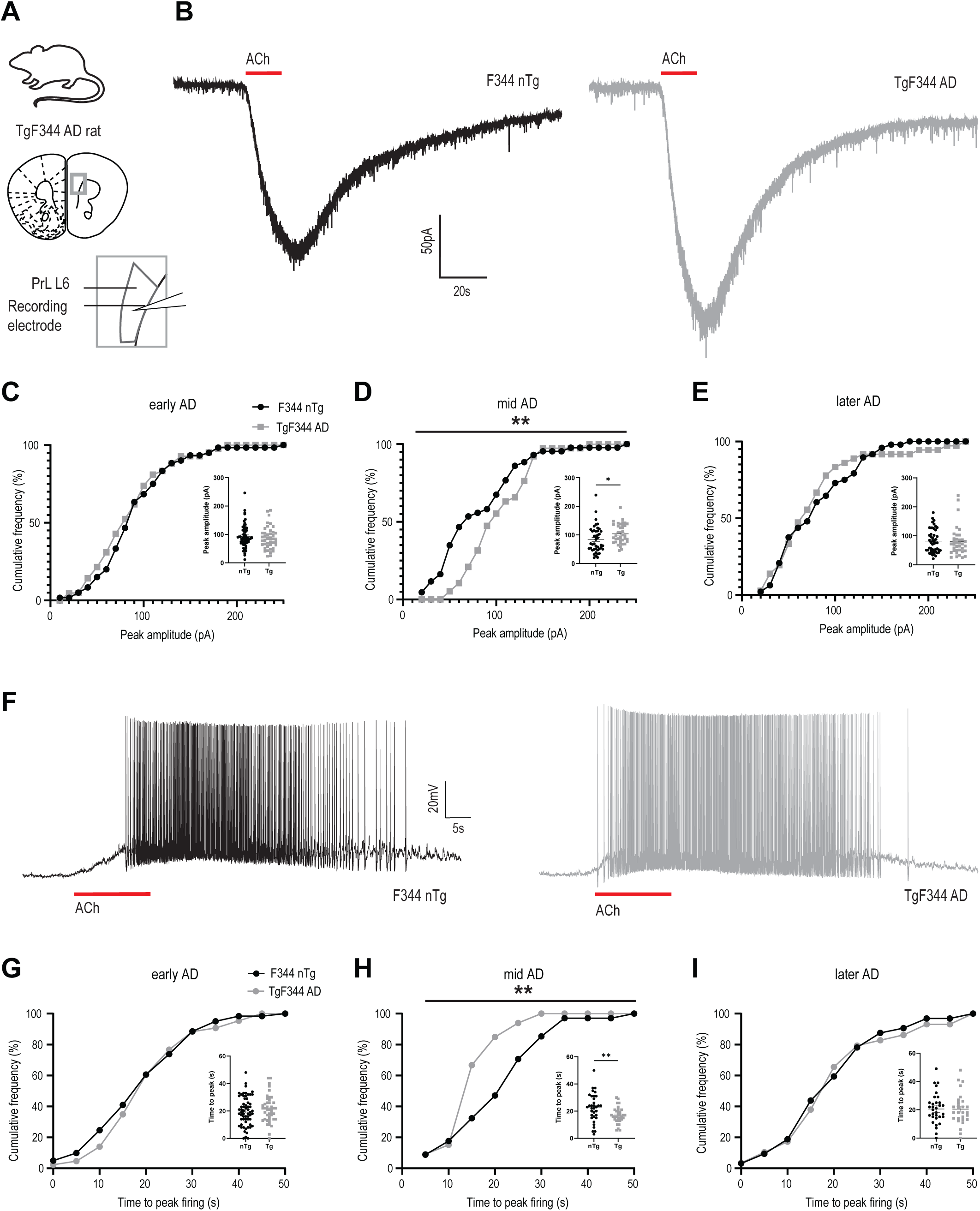
Early cholinergic upregulation conserved in different species and AD model. **A** Image depicts TgF344 AD rat model and coronal brain section with the recording electrode positioned over layer 6 of the prefrontal cortex. **B** Example responses to exogenous bath-applied acetylcholine from F344 non-transgenic (nTg) control (black) and TgF344 AD (grey) neurons measured in voltage clamp (Vm = –75 mV). Graphs show cumulative frequency of peak amplitude of acetylcholine current is **C** not different at 8 months, **D** is significantly higher for TgF344 AD responses than controls at 12 months (*P* < 0.01), **E** and is not different at 18 months. Insets show the data in scattergram. **F** Example responses to exogenous bath-applied acetylcholine from F344 WT control (black) and TgF344 AD (grey) neurons measured in current clamp. Graphs show cumulative frequency of time to peak firing of acetylcholine response is **G** not different at 8 months, **H** is significantly faster for TgF344 AD responses than controls at 12 months (*P* < 0.01), **I** and is not different at 18 months. Insets show the data in scattergram. Rats aged in three cohorts: 8 months (nTg: 8.5 ± 0.2 months, n = 12, TgF344: 8.4 ± 0.2, n = 9), 12 months (nTg: 12.4 ± 0.2 months, n = 13, TgF344 12.5 ± 0.2 months, n = 10), and 18 months (nTg: 17.7 ± 0.3 months, n = 14, TgF344: 17.6 ± 0.2 months, n = 12).

A similar pattern is observed for changes in action potential firing in response to cholinergic stimulation (Fig 3F). The TgF344-AD rat model shows an upregulation in cholinergic firing, as measured by the reduced time taken to attain peak firing rate relative to controls at mid-AD (Fig 3G-I, KS test, *P* = 0.003, D = 0.4). However, this difference was not observed at early-AD (KS test, *P* = 0.8, D = 0.1) nor at later-AD (KS test, *P* = 1, D = 0.1).

To delve deeper into the cholinergic responses, we examine the impact of sex within each age group. We detect no sex differences in ACh currents (two-way ANOVA, 8 months: F_1, 98_ = 0.4, *P* = 0.5; 12 months: F_1, 77_ = 0.04, *P* = 0.8; 18 months: F_1, 80_ = 0.003, *P* = 0.9) nor significant interactions between sex and genotype (8 months: F_1, 98_ = 2.7, *P* = 0.1; 12 months: F_1, 77_ = 0.09, *P* = 0.8; 18 months: F_1, 80_ = 0.3, *P* = 0.6).

The differences in cholinergic responses observed in the mid-AD (12 month) group occur in the absence of corresponding differences in intrinsic properties (Supplemental Table 2). Examination of the excitability of these neurons with input-output analysis (Fig S2A-C) shows a significant decrease in the intrinsic excitability curve of TgF344 AD neurons at 12 months (Fig S2B; nonlinear regression, comparison of fit, F_3,561_ = 3.8, *P* = 0.01). Interestingly, this decrease in excitability to current injection coincides with the emergence of increased excitability to cholinergic stimulation (Fig 3). This finding underscores the relative specificity of increased cholinergic excitation of layer 6 neurons at mid-AD in this rat model. In the early AD (8 month) and later-AD (18 month) groups, we observe no significant differences in layer 6 neuronal properties (Supplemental Table S2) nor intrinsic excitability (Fig S2A,C; F test = ns).

Together, this work shows the replication of the AD upregulation in cholinergic responses in layer 6 pyramidal neurons in a different species, a different model, and with a different method of cholinergic stimulation. This conserved property underscores the importance of understanding the receptor mechanisms responsible.

### Upregulated responses arise from a selectively-increased nicotinic signal

Since we find that AD cholinergic responses are faster and greater amplitude than non-transgenic controls, we hypothesize that increased nicotinic responses underlie the genotype difference, as these ionotropic receptors impart these characteristics to the endogenous response [62,63,67]. To investigate pre-and post-synaptic elements in the intact prefrontal cholinergic synapse, we use the opto-TgCRND8 AD mouse. To investigate the nicotinic receptor contribution to this upregulated opto-ACh response in AD, we pharmacologically block these receptors using the antagonist DHβE (3 µM) (Fig 4A). DHβE changes the cholinergic responses in both genotypes (Fig 4B), significantly decreasing the rise speed (paired, two-tailed t-test: *t* = 4.3, *P* = 0.0002, df = 23) and the peak amplitude (paired, two-tailed t-test: *t* = 3.5, *P* = 0.002, df = 23). However, DHβE exerts a greater effect on the AD group, yielding a significantly greater change in rise slope (Fig 4C, unpaired, two-tailed *t*-test, *t* = 2.3, *P* = 0.03, df = 22) and peak amplitude (Fig 4D, unpaired, two-tailed *t*-test, *t* = 2.6, *P* = 0.01, df = 22). These results suggest that increased opto-ACh responses in this preclinical AD model include a prominent upregulation in nicotinic receptor signalling.

**Figure 4.**
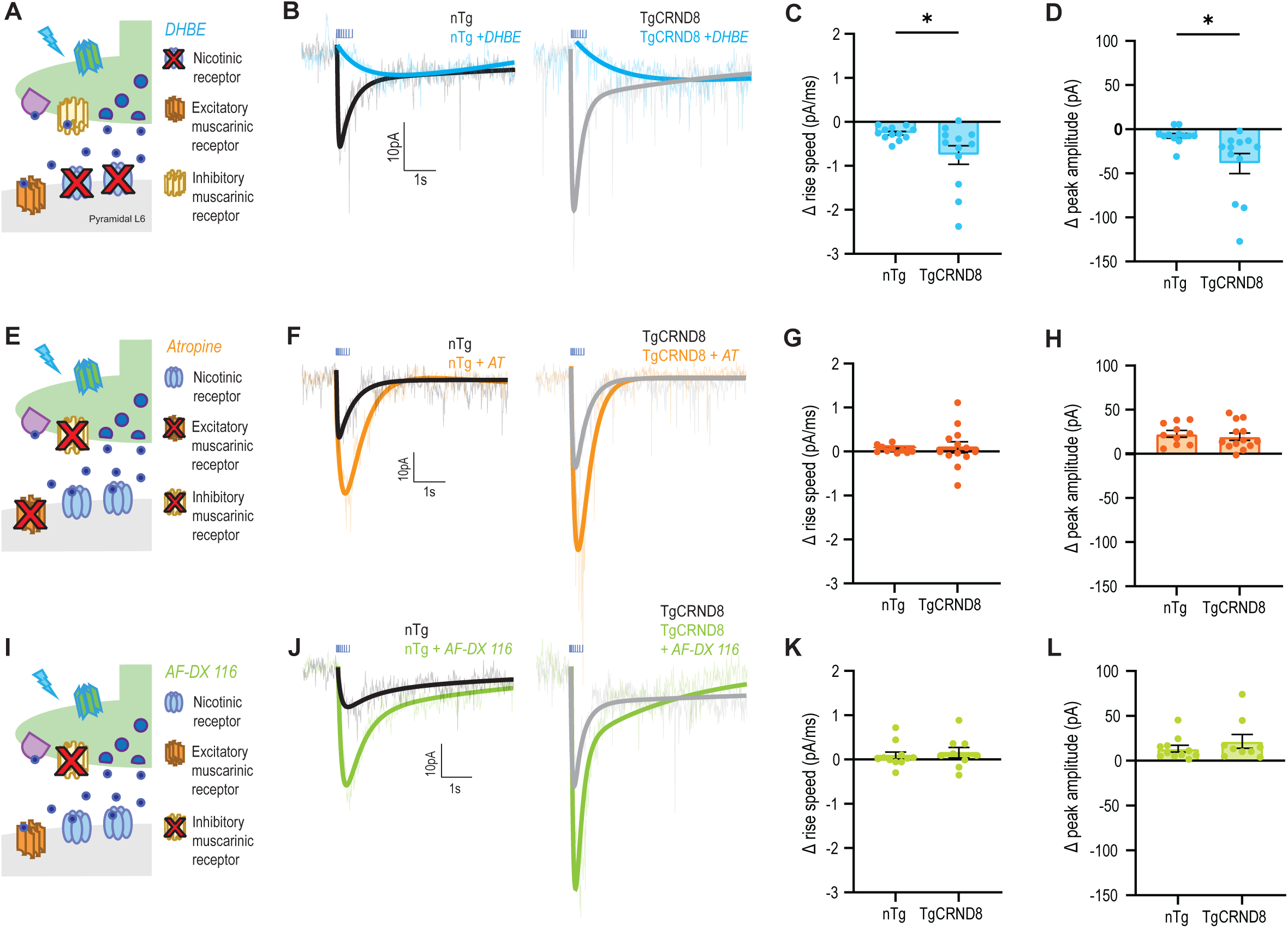
Early cholinergic upregulation in AD model is specific to nicotinic receptor signalling. **A** Schematic depicts cholinergic synapse in the presence of nicotinic receptor antagonist DHβE. **B** Paired example opto-ACh responses from non-transgenic (nTg) control (black) and TgCRND8 AD (grey) neurons measured in voltage clamp (Vm = –75mV) before and after DHβE. Graphs show that DHβE causes a greater decline in **C** rise speed (*P* < 0.05) and **D** peak amplitude (*P* < 0.05) of opto-ACh currents in TgCRND8 AD responses relative to controls (age range: 4.5 to 12.2 months, mean = 7.9 ± 0.7 months, n = 14). **E** Schematic depicts cholinergic synapse in the presence of broad muscarinic receptor antagonist atropine. **F** Paired example opto-ACh responses from non-transgenic (nTg) control (black) and TgCRND8 AD (grey) neurons measured in voltage clamp before and after atropine. Graphs show that atropine elicits changes in **G** rise speed and **H** peak amplitude that are not different between TgCRND8 AD and control responses (age range: 3.1 to 12.2 months, mean = 6.4 ± 0.7 months, n = 20). **I** Schematic depicts cholinergic synapse in the presence of muscarinic M2 receptor antagonist AF-DX 116. **J** Paired example opto-ACh responses from non-transgenic (nTg) control (black) and TgCRND8 AD (grey) neurons measured in voltage clamp before and after AF-DX 116. Graphs show that AF-DX 116 elicits changes in **K** rise speed and **L** peak amplitude that are not different between TgCRND8 AD and control responses (age range: 4.2 to 12.2 months, mean = 7.8 ± 0.9 months, n = 13).

Since G-protein coupled muscarinic receptors contribute both pre– and post-synaptically to prefrontal cholinergic synapses [62], we next evaluate whether muscarinic components of the cholinergic synapse are altered. Blocking muscarinic receptors with the broad-spectrum antagonist, atropine (200nM; Fig 4E), increases the amplitude of cholinergic responses in both genotypes (Fig 4F; paired, two-tailed *t*-test: *t* = 7.4, *P* < 0.0001, df = 22), but does not change the rise slope (paired, two-tailed *t*-test: *t* = 1.1, *P* = 0.3, df = 22). However, there is no significant genotype difference in the effect of atropine on either of these measures (Fig 4G, rise slope: unpaired, two-tailed *t*-test: *t* = 0.3, *P* = 0.8, df = 21; Fig 4H, peak amplitude: unpaired, two-tailed *t*-test: *t* = 0.5, *P* = 0.6, df = 21). Consistent with our interpretation that atropine increases opto-ACh response amplitude by blocking presynaptic autoinhibition, targeted block of presynaptic muscarinic autoreceptors with AF-DX 116 (Fig 4I) also increases opto-ACh responses in both AD and control (Fig 4J). AF-DX 116 increases opto-ACh response amplitude (paired, two-tailed *t*-test: *t* = 4.2, *P* = 0.005, df = 19) but does not change the rise slope (paired, two-tailed *t*-test: *t* = 1.7, *P* = 0.1, df = 19). There is no significant genotype difference in the effect of AF-DX 116 on either of these measures (Fig 4K, rise slope: unpaired, two-tailed *t*-test: *t* = 0.5, *P* = 0.7, df = 18; Fig 4L, peak amplitude: unpaired two-tailed *t*-test: *t* = 1, *P* = 0.3, df = 18). These findings suggest that neither pre-nor post-synaptic muscarinic receptors are substantially altered in this preclinical AD model.

### Treatment strategies: acetylcholinesterase inhibition and nicotinic potentiation amplify cholinergic upregulation

To probe the impact of treatment intervention on this upregulated cholinergic signalling, we measure the effects of current standard of care treatments and a novel treatment strategy on the amplitude and kinetics of the AD neuron and non-transgenic control opto-ACh responses from early to mid and later-disease in the TgCRND8 AD mouse.

The pro-cognitive standard-of-care treatment galantamine inhibits acetylcholinesterase (Fig 5A), allowing acetylcholine to remain longer at the synapse. Applying galantamine to opto-ACh AD responses (Fig 5B) trends towards increasing the rise speed (Fig 5C, paired, two-tailed *t*-test, *t* = 2.3, *P* = 0.06, df = 7) and significantly increases peak amplitude (Fig 5D, paired, two-tailed *t*-test, *t* = 4.4, *P* = 0.003, df = 7). However, its greatest impact is to significantly prolong the decay constant (Fig 5E, paired, two-tailed *t*-test, *t* = 3.6, *P* = 0.008, df = 7). This effect of galantamine is consistent across genotype, in TgCRND8 mice and non-transgenic controls (Fig S3, A-C) and consistent in TgCRND8 mice at early to mid-AD as well as late AD (Fig S4, A-C). Galantamine modestly increases the peak response magnitude; however, it prolongs the overall kinetics of the usually brief cholinergic response.

**Figure 5.**
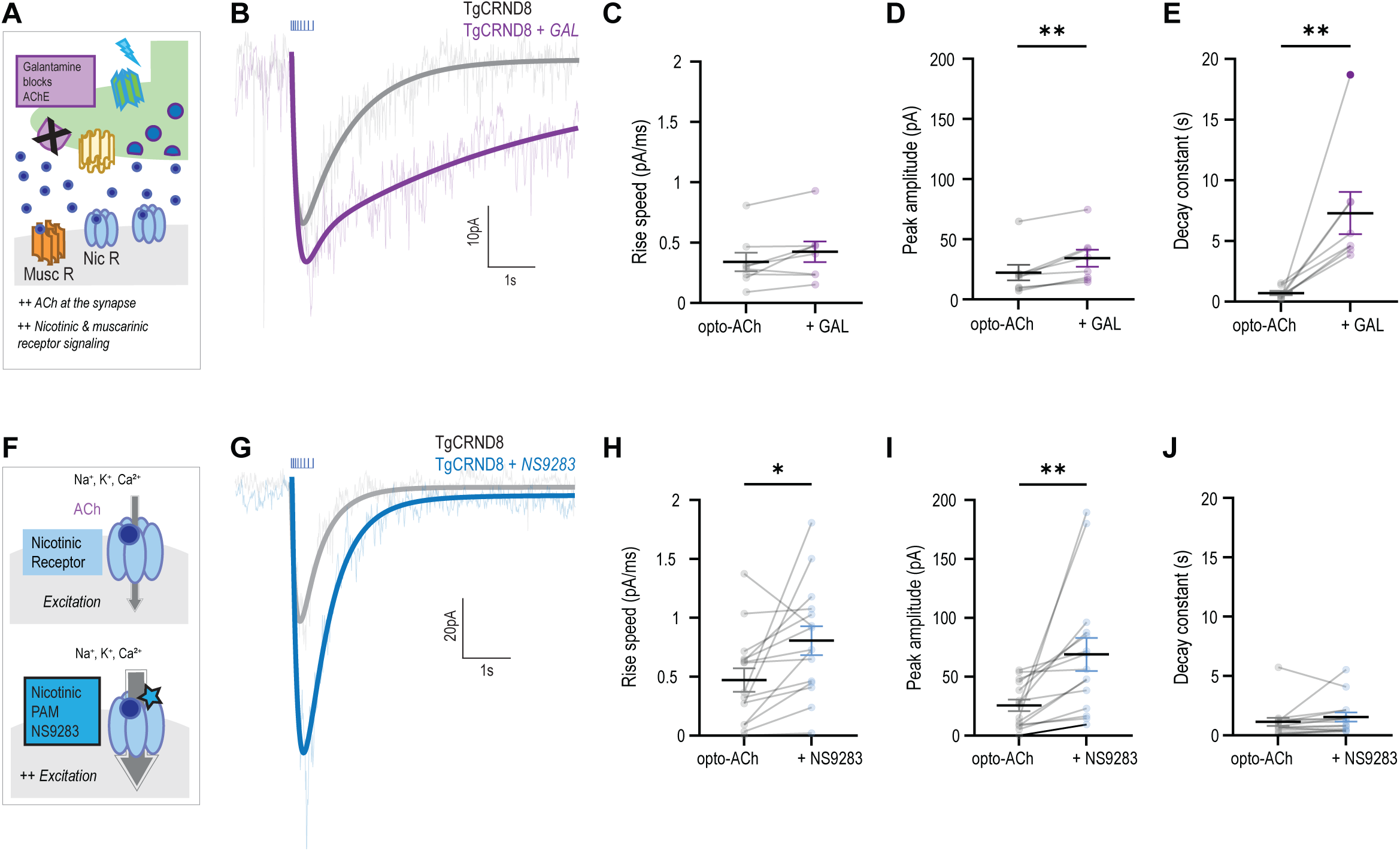
Galantamine and a novel nicotinic treatment strategy further enhances cholinergic upregulation in AD model. **A** Schematic depicts cholinergic synapse in the presence of acetylcholinesterase inhibitor galantamine. Galantamine slows the degradation of acetylcholine, prolonging its availability at the synapse. **B** Paired example opto-ACh response from a TgCRND8 AD neuron before (grey) and after galantamine (purple). Graphs show that galantamine elicits **C** no change in rise speed but a significant increase in **D** peak amplitude (*P* < 0.01) and **E** response duration, as measured by the decay constant (*P* < 0.01) in TgCRND8 AD responses (age range 3-6 months: mean = 5.2 ± 0.6 months, n = 5). **F** Schematic depicts the activation of postsynaptic nicotinic receptors by acetylcholine and the action of nicotinic PAM NS9283 in increasing nicotinic receptor conductance. **G** Paired example opto-ACh responses from a TgCRND8 AD neuron before (grey) and after NS9283 (blue). Graphs show that NS9283 elicits an increase in **H** rise speed (*P* < 0.05) and **I** peak amplitude (*P* < 0.001) **J** with no change in the response duration (age range 3-6 months: mean = 4.7 ± 0.4 months, n = 6).

To explore treatment interventions targeted to the specific nicotinic receptor upregulation seen in this AD model, we test the nicotinic-selective positive allosteric modulator (PAM), NS9283 to enhance endogenous cholinergic signalling (Fig 5F). We find NS9283 greatly boosts opto-ACh AD responses (Fig 5G), significantly increasing the rise speed (Fig 5H; paired, two-tailed *t*-test, *t* = 2.7, *P* = 0.02, df = 14) and the peak amplitude (Fig 5I; paired, two-tailed *t*-test, *t* = 3.2, *P* = 0.007, df = 14). Strikingly, these changes occur without changing the decay constant (Fig 5J; paired, two-tailed *t*-test, *t* = 1.3, *P* = 0.2, df = 14) of the opto-ACh response. The action of NS9283 is consistent across genotype, in TgCRND8 mice and non-transgenic controls (Fig S3, D-F) and consistent in TgCRND8 mice at early to mid-AD as well as late AD (Fig S4, D-F). This intervention strongly amplifies peak response magnitude while retaining the typical rapid and brief response kinetics.

The current standard of care treatment galantamine significantly increases response amplitude (71% ± 16%) and trends towards a significant increase in rise speed (35% ± 16%), but at the cost of significantly slowed decay of the cholinergic response (2032% ± 993%). In comparison, the nicotinic specific NS9283 achieves a significant enhancement in rise speed (210% ± 68%), a significant increase in amplitude (249% ± 87%) and modestly but not consistently prolongs response decay (77% ± 32%). NS9283 incites a greater increase in response magnitude and better conserves endogenous response timing and kinetics than the broad cholinergic standard of care treatment. We demonstrate it is possible to use a targeted treatment such as NS9283 to further enhance upregulation of this cognitively-crucial signalling pathway.

## Discussion

We demonstrate a functional upregulation in layer 6 prefrontal cholinergic signalling in preclinical models of AD. Cholinergic upregulation occurs during early to mid-disease progression in two species models with differing pathology trajectories. Elevated functional nicotinic signalling plays a dominant role, and the apparent compensation is sensitive to further enhancement via pharmacological manipulation. The AD treatment of acetylcholinesterase inhibition boosts the cholinergic response but slows its time course. By contrast, targeted nicotinic positive allosteric modulation exerts a strong and significant potentiation with preserved kinetics. Taken together, our results suggest a potential active mechanism of cognitive compensation and demonstrate an effective approach to amplify and potentially extend it further into the disease process.

### Cognitive reserve and neural compensation in AD

Cognitive reserve, or the brain’s ability to sustain neurological decline while maintaining cognitive processes, is well documented clinically in aging [12,13,70,71] and in AD [72,73]. Emerging work explores the neurophysiological basis for cognitive resilience in AD [15,74] including in the prefrontal cortex [75]. In preclinical models, plasticity related to AD has been detected in cortical neuronal morphology [76–83] and in neurotransmitter signalling [84,85], including acetylcholine neuromodulation [86]. In the TgF344 AD rat model used here, for example, parvalbumin interneuron neuroplasticity is observed mid-pathology, with coinciding cognitive resilience in executive function and cognitive flexibility [14]. Interestingly, this emergence of neuroplasticity and cognitive resilience is observed at the same mid-disease timepoint as the cholinergic upregulation in layer 6 found here.

Prefrontal cholinergic signalling is crucial for many cognitive functions maintained by cognitive reserve, including attention, cue detection, and working memory [18–20,22,23]. The nicotinic excitation of layer 6 pyramidal neurons is thought to play a key role in attention and cue detection [21,24]. The importance of prefrontal nicotinic signalling in successful attention [87–90] becomes evident under demanding conditions, where locally stimulating nicotinic receptors improves performance [91] and deficient prefrontal nicotinic signalling worsens performance [21,24,92]. These cognitive processes are timescale dependent [93–95], where configurations of the fast ionotropic nicotinic receptor that include the α5 subunit lead to faster responses to endogenous acetylcholine [63] and contribute to better attentional performance [21]. The nicotinic positive allosteric modulator NS9283, which potentiates cholinergic responses while maintaining fast endogenous kinetics, improves performance on attention tasks in wildtype rodents [96,97].

Clinically, in individuals pre-diagnosis with elevated amyloid-β and none/mild baseline cognitive deficits, cholinergic inhibition unmasks otherwise undetectable cognitive deficits that correlate with amyloid-β load [98]. This supports that in pre-diagnosis or early AD, cholinergic signalling may be important in compensating for pathological decline and maintaining cognitive functions. This preclinical and clinical behavioral literature, coupled with the opto-physiological data in the current manuscript, underscores the need for future work to examine the behavioral impact of nicotinic-targeted therapeutics such as NS9283 in models of AD and, particularly, its interactions with age and stage of disease.

### Cholinergic signalling and synapses in AD

Previous molecular investigations into cholinergic neurons and synapses in AD emphasize a trajectory of cholinergic decline, with decreases in both pre– and post-synaptic markers resulting in eventual heterogenous loss of cholinergic neurons in advanced AD [32,33,99–109]. However, much of this characterization is performed later in disease progression [110,111]. Molecular and functional investigation that has been carried out in early to mid-disease mostly consists of swiftly-progressing genetic models of AD that preclude the use of littermate controls [38–44]. To detect and systematically investigate mechanisms of compensatory plasticity, it is essential to use a well-charted model with appropriately-matched non-transgenic controls. Furthermore, functional characterisation is vital for understanding cholinergic synaptic transmission.

Here, we find an upregulation in prefrontal cholinergic signalling in layer 6 that emerges after disease onset, at early to mid-disease in two different preclinical models of AD. Both the TgCRND8 AD mouse and the TgF344 AD rat used here employ human APP mutations [48,57], resulting in amyloid-driven AD pathologies. Amyloid-β interacts with nicotinic receptors [112–119], the consequences of which depend on cell type and nicotinic receptor configuration. Nicotinic receptors are regulated by many cellular elements, including lynx family proteins [67,120] and protein kinase C by way of muscarinic receptor agonism [62], providing many potential avenues for signalling to be harnessed for compensation. Our use of models with littermate non-transgenic controls reveals an upregulation of nicotinic signalling in early to mid-AD that does not continue to increase in later-AD. Strengths of our electrophysiological and opto-physiological approach include functional insight into cholinergic signalling upregulation and identification of this upregulation as specific to nicotinic receptors.

A caveat for our work is its limitation to layer 6 pyramidal neurons. While this neuronal population has prominent expression of nicotinic receptors and their regulatory lynx prototoxins [67] and plays a key role in attention [21,24,92], processes of cognitive reserve likely also require the involvement of neurons in other prefrontal layers. It will be important for future work to specifically target the impact of endogenous cholinergic responses on additional neurophysiological processes beyond those involved in attention.

### Translational and clinical impact

Deficits in attention and executive function have consequences for other cognitive domains: if you can’t attend, you can’t encode [121]. It is therefore important to identify mechanisms through which we can improve attention and extend the maintenance of this cognitive domain as much as possible through AD progression. Nicotinic signalling has long been a target for enhancing cognition, including in AD [122,123]. Direct stimulation of nicotinic receptors is pro-cognitive in AD [27], potentially protective against amyloid beta [124,125] but complicated by desensitization [126–128]. Nicotinic positive allosteric modulators, such as NS9283 used here, present less risk of desensitization [129], and are pro-cognitive preclinically [96,97]. Our results suggest that nicotinic positive allosteric modulation further enhances nicotinic signalling upregulation in AD.

Attention and cue-detection are mediated by cholinergic, and specifically nicotinic stimulation of the prefrontal cortex [19,20,87–90,130] and depend on fast ionotropic receptor kinetics [21,63]. Acetylcholinesterase inhibitor, a current and long-used AD treatment, enhances all cholinergic signaling: fast, ionotropic nicotinic signal and slower, longer-lasting metabotropic muscarinic signal shown above (Figure 5) and previously [62,63]. While acetylcholinesterase inhibitor imparts some increase in response amplitude, it profoundly slows response decay. Treatments that maintain the endogenous timing of responses, such as NS9283 that increases the pro-attentional rise kinetics, largely amplifies response magnitude, and does not alter decay might better enhance time-sensitive cognitive functions.

Understanding neural compensation and neurological changes at different disease stages is important for understanding the implications of clinical interventions. Differing levels of effectiveness of novel AD treatments may result from timing of intervention and the level of neurological changes already set in motion by AD. Adaptations that are pro-cognitive in the presence of AD may have altered outcomes if pathology is resolved. Therefore, it is important to take neurological changes into account when determining treatment strategies.

## Conclusion

Overall, we uncover evidence for cholinergic compensation in two preclinical models of AD. In both models, functional upregulation in prefrontal cholinergic signalling occurs in early to mid-AD. Mechanistic investigation reveals increased nicotinic signalling, which is amenable to further upregulation, both by acetylcholinesterase inhibitor and by nicotinic positive allosteric modulation. While the scope of benefit across cognitive domains requires investigation, harnessing compensatory nicotinic upregulation may be an important treatment strategy to further enhance and preserve cognition in AD.

## Declarations

### Contributions

Conceptualization, SKP, JM, EKL; Methodology, SKP, SV, EKL; Data acquisition SKP, SV; Analysis, SKP, SQ; Writing, SKP EKL; Review & editing, SKP, SV, JM, EKL; Funding, SKP, JM, EKL; Supervision, JM, EKL. All authors have critically reviewed and approved of the paper.

## Abbreviations

ACh: acetylcholine
AD: Alzheimer’s Disease
ChAT: choline acetyl transferase
ChR2: channelrhodopsin
DHßE: dihydro-ß-erythroidine
gal: galantamine
hz: hertz
ms: millisecond
mV: millivolt
nTg: non-transgenic
opto-Ach: optogenetic release of acetylcholine
pA: pico-ampere
PAM: positive allosteric modulator
prl: prelimbic
Tg: transgenic

## Acknowledgements

We thank Janice McNabb and Yao-Fang Tan for expert technical assistance, and Madeleine Ridgeway for optimizing analysis of a related dataset.

## Ethics approval – animals

This manuscript contains animal research that was approved by the the Faculty of Medicine Animal Care Committee at the University of Toronto in accordance with the guidelines of the Canadian Council on Animal Care (protocols #20011123, #20011621, #20011796). This work was conducted under the appropriate Material Transfer Authorizations

## Ethics approval – humans

n/a

## Consent for publication

n/a

## Availability of data

The datasets generated during and/or analysed during the current study are not publicly available due to agency over intellectual property but are available from the corresponding author on reasonable request.

