## Supplemental Materials for "Enhanced prefrontal nicotinic signaling as evidence of active compensation in Alzheimer’s disease models"

- Supplemental Tables S1 & S2
- Supplemental Figures S1, S2, S3, & S4

**Supplemental Table S1**

| | Capacitance | Input resistance | Resting membrane potential | Threshold | Spike amplitude |
|---|---|---|---|---|---|
| nTg<br>3-6 months | 85 ± 2 pF | 137 ± 5 MΩ | -85 ± 1 mV | -50 ± 1 mV | 63 ± 3 mV |
| TgCRND8<br>3-6 months | 87 ± 3 pF | 127 ± 6 MΩ | -84 ± 1 mV | -50 ± 1 mV | 69 ± 2 mV |
| Statistics | $t_{(112)} = 0.5$<br>$P = 0.6$ | $t_{(112)} = 1.1$<br>$P = 0.3$ | $t_{(112)} = 0.6$<br>$P = 0.6$ | $t_{(112)} = 0.5$<br>$P = 0.6$ | $t_{(112)} = 1.8$<br>$P = 0.08$ |
| Summary | *NS* | *NS* | *NS* | *NS* | *NS* |
| nTg<br>7-10 months | 87 ± 1 pF | 143 ± 6 MΩ | -80 ± 3 mV | -50 ± 1 mV | 67 ± 2 mV |
| TgCRND8<br>7-10 months | 89 ± 2 pF | 160 ± 9 MΩ | -82 ± 2 mV | -48 ± 1 mV | 65 ± 3 mV |
| Statistics | $t_{(99)} = 0.7$<br>$P = 0.5$ | $t_{(99)} = 1.5$<br>$P = 0.1$ | $t_{(99)} = 0.6$<br>$P = 0.6$ | $t_{(99)} = 1.2$<br>$P = 0.2$ | $t_{(99)} = 0.7$<br>$P = 0.5$ |
| Summary | *NS* | *NS* | *NS* | *NS* | *NS* |

Neuronal intrinsic properties of layer 6 pyramidal neurons by genotype in mouse. Table shows mean ± SEM for each intrinsic property in neurons from non-transgenic (nTg) and TgCRND8 AD model mice at early/mid AD (3 to 6 months) and later AD (7-10 months), as well as the results of the unpaired *t*-tests comparing genotypes in each age group.

**Supplemental Table S2**

| | Capacitance | Input resistance | Resting membrane potential | Threshold | Spike amplitude |
|---|---|---|---|---|---|
| F344 nTg<br>8 months | 104 ± 4 pF | 96 ± 4 MΩ | -91 ± 1 mV | -52 ± 1 | 72 ± 1 mV |
| TgF344<br>8 months | 96 ± 4 pF | 101 ± 5 MΩ | -91 ± 1 mV | -52 ± 1 | 69 ± 1 mV |
| Statistics | $t_{(107)}$ = 1.5<br>*P* = 0.1 | $t_{(107)}$ = 0.4<br>*P* = 0.7 | $t_{(107)}$ = 0.5<br>*P* = 0.6 | $t_{(107)}$ = 0.05<br>*P* = 0.9 | $t_{(107)}$ = 1.7<br>*P* = 0.9 |
| Summary | *NS* | *NS* | *NS* | *NS* | *NS* |
| F344 nTg<br>12 months | 101 ± 4 pF | 104 ± 7 MΩ | -89 ± 1 mV | -51 ± 1 | 72 ± 2 mV |
| TgF344<br>12 months | 95 ± 3.5 pF | 87 ± 4 MΩ | -90 ± 1 mV | -50 ± 1 | 67 ± 2 mV |
| Statistics | $t_{(85)}$ = 0.5<br>*P* = 0.6 | $t_{(85)}$ = 1.9<br>*P* = 0.06 | $t_{(85)}$ = 0.2<br>*P* = 0.9 | $t_{(85)}$ = 1.3<br>*P* = 0.2 | $t_{(85)}$ = 1.8<br>*P* = 0.08 |
| Summary | *NS* | *NS* | *NS* | *NS* | *NS* |
| F344 nTg<br>18 months | 105 ± 4 pF | 112 ± 6 MΩ | -88 ± 1 mV | -49 ± 1 | 67 ± 2 mV |
| TgF344<br>18 months | 95 ± 4 pF | 127 ± 8 MΩ | -88 ± 1 mV | -50 ± 1 | 69 ± 1 mV |
| Statistics | $t_{(110)}$ = 1.8<br>*P* = 0.07 | $t_{(110)}$ = 1.5<br>*P* = 0.1 | $t_{(110)}$ = 0.4<br>*P* = 0.7 | $t_{(110)}$ = 1.4<br>*P* = 0.1 | $t_{(110)}$ = 1.4<br>*P* = 0.2 |
| Summary | *NS* | *NS* | *NS* | *NS* | *NS* |

Neuronal intrinsic properties of layer 6 pyramidal neurons by genotype in rat. Table shows mean ± SEM for each intrinsic property in neurons from F344 non-transgenic (nTg) and TgF344 AD model rats at each age point, as well as the results of the unpaired *t*-tests comparing genotypes at each age point.

### Supplemental Figure S1

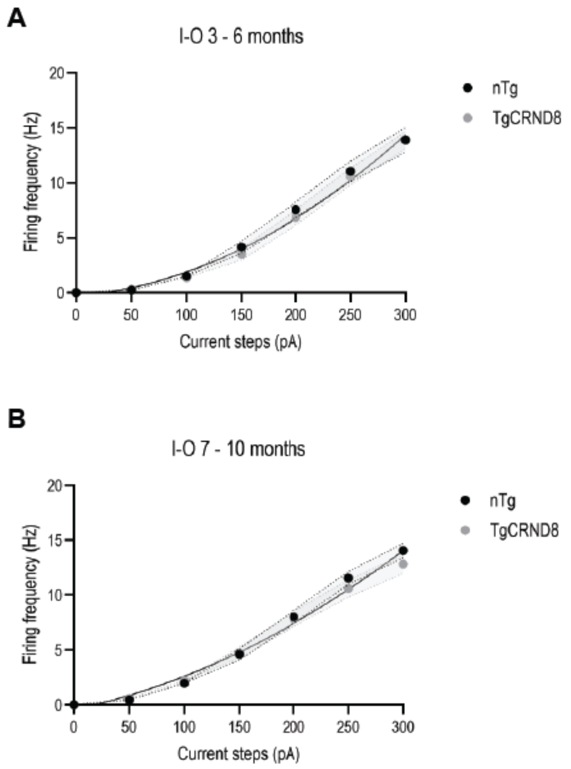


**Intrinsic excitability comparison between nTg and TgCRND8 layer 6 pyramidal neurons.** Input-output graphs show the firing frequency of pyramidal layer 6 prefrontal cortex neurons in response to increasing steps of injected current in **A** neurons from 3 to 6 month old TgCRND8 and nTg animals and **B** neurons from 7 to 10 month old TgCRND8 and nTg animals. There is no significant difference between intrinsic excitability of neurons of the younger (nonlinear regression, comparison of fit, $F_{3,750}$ = 0.4, $P$ = 0.7) nor older age group ($F_{3,788}$ = 1.6, $P$ = 0.1). Shaded area shows SEM.

**Supplemental Figure S2**

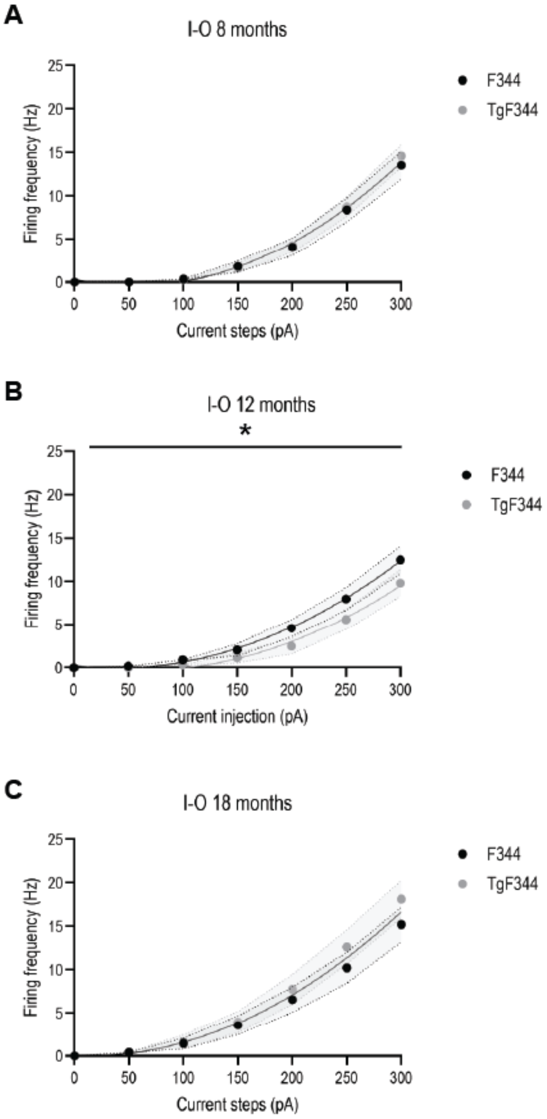


**Intrinsic excitability comparison between F344 and TgF344 layer 6 pyramidal neurons.** Input-output graphs show the firing frequency of pyramidal layer 6 prefrontal cortex neurons in response to increasing steps of injected current in neurons from TgF344 and F344 control animals at **A** 8 month old, **B** 12 month old, and **C** 18 month old. At 12 months, there is a small but significant decrease in intrinsic excitability of TgF344 AD neurons (nonlinear regression, comparison of fit, $F_{3,561} = 3.8$, $P = 0.011$). This coincides with the emergence of increased cholinergic excitability (see **Figure 3**). There are no significant differences in intrinsic excitability of pyramidal L6 neurons at 8 months ($F_{3,603} = 0.3$ $P = 0.8$) nor 18 months, ($F_{3,449} = 1.5$, $P = 0.2$). Shaded area shows SEM.

**Supplemental Figure S3**

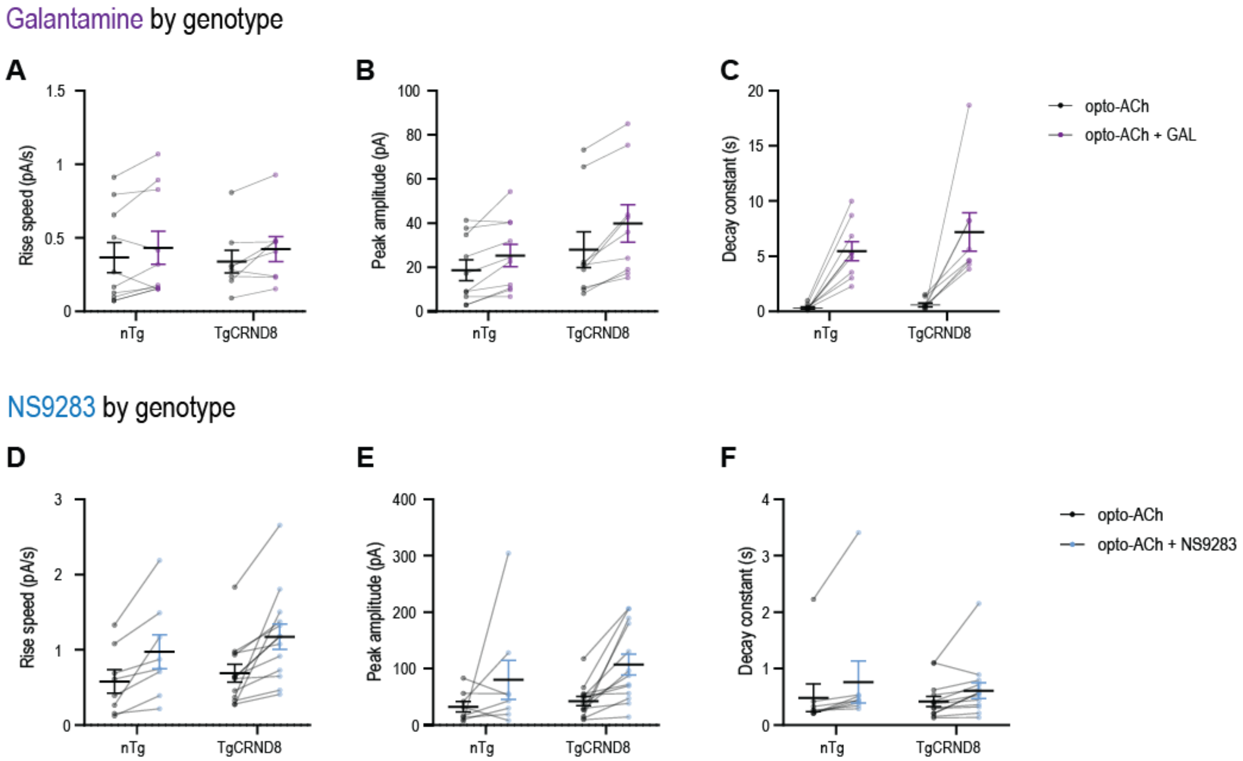


**Cholinergic treatments are consistently effective in prefrontal layer 6 pyramidal neurons of TgCRND8 AD and nTg controls.** Graphs show the effect of galantamine (purple) and NS9283 (blue) on opto-ACh responses in non-transgenic controls (nTg) as compared to TgCRND8 mice. Galantamine elicits no significant effect on **A** rise speed or **B** peak amplitude in either genotype but does elicit a significant drug effect on **C** decay constant ($P < 0.0001$, drug effect, two-way ANOVA) (nTg: 4 ± 0.4 months, n = 6; TgCRND8: 5.2 ± 0.6 months, n = 5). NS9283 elicits a significant drug effect on **D** rise speed ($P = 0.01$, drug effect, two-way ANOVA) and **E** peak amplitude ($P = 0.006$, drug effect, two-way ANOVA) for both genotypes with no significant effect on **F** decay (nTg: 4.1 ± 0.7 months, n = 5; TgCRND8: 4.7 ± 0.4 months, n = 6). There are no interactions between genotype and drug treatment. Graphs show mean ± SEM.

**Supplemental Figure S4**

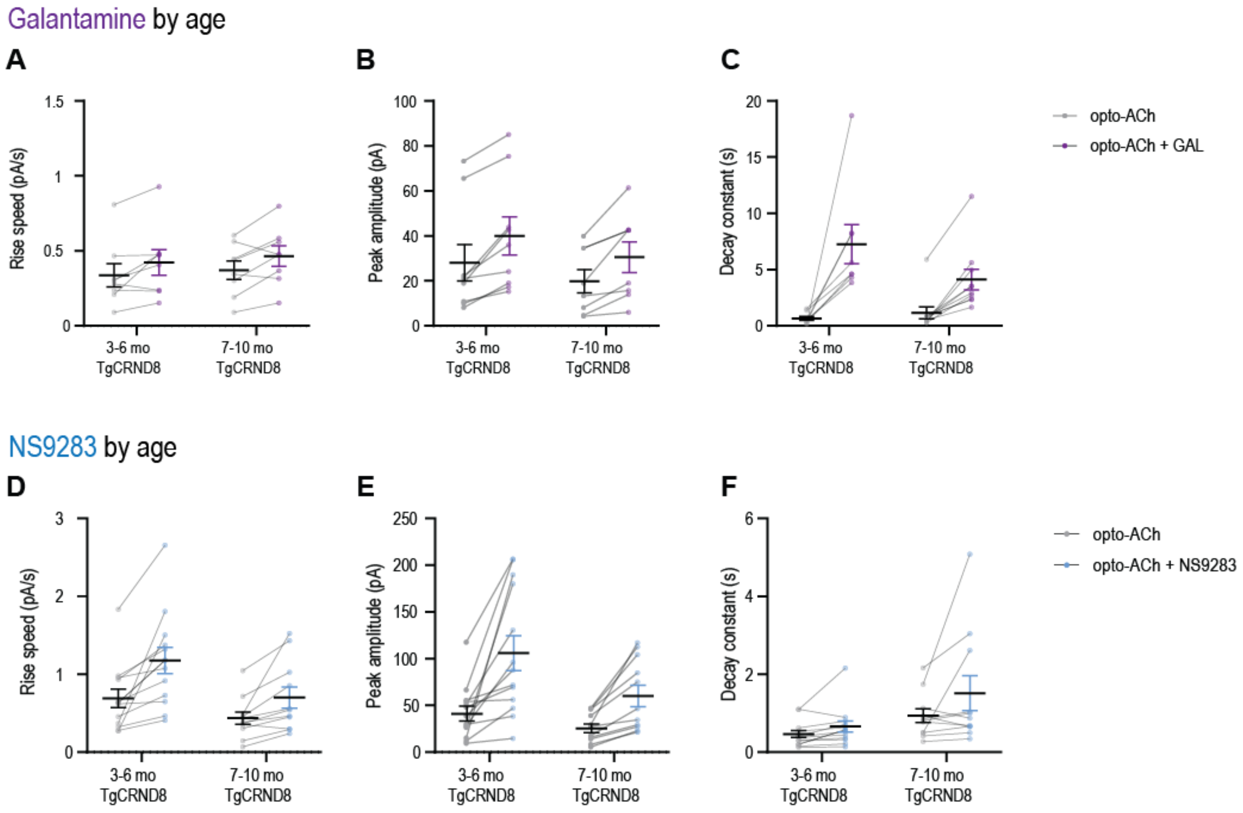


**Cholinergic treatments are consistently effective at early to mid-AD and late-AD in layer 6 pyramidal neurons of prefrontal cortex.** Graphs show the effect of galantamine (purple) and NS9283 (blue) on opto-ACh responses in early to mid-AD TgCRND8 mice (3-6 months) as compared to late AD TgCRND8 mice (7-12 months). Galantamine elicits no significant drug effect on **A** rise speed or **B** peak amplitude for either age point but elicits a significant drug effect on **C** decay (*P* < 0.0001, drug effect, two-way ANOVA) (3-6 months: 5.2 ± 0.6 months, n = 5; 7-12 months: 8.5 ± 0.3, n = 4). NS9283 elicits a significant effect on **D** rise speed (*P* = 0.0007, drug effect, *P* = 0.0009, age effect, two-way ANOVA) and **E** peak amplitude (*P* = 0.0003, drug effect, *P* = 0.02, age effect, two-way ANOVA) for both age points with no significant effect on **F** decay (3-6 months: 4.7 ± 0.4 months, n = 6; 7-12 months: 8.3 ± 0.7 months, n = 5). There are no interactions between age and drug treatment. Graphs show mean ± SEM.
